# Glucose metabolism promotes neonatal heart regeneration

**DOI:** 10.1101/865790

**Authors:** Viviana M Fajardo Martinez, Iris Feng, Bao Ying Chen, Cesar A Perez, Baochen Shi, Peter Clark, Rong Tian, Ching-Ling Lien, Matteo Pellegrini, Heather Christofk, Haruko Nakano, Atsushi Nakano

**Affiliations:** Division of Pediatrics, David Geffen School of Medicine, University of California, Los Angeles; Department of Molecular Cell Developmental Biology, School of Life Science, University of California, Los Angeles; Department of Molecular and Medical Pharmacology, University of California, Los Angeles; Department of Chemistry and Biochemistry, University of California, Los Angeles; Crump Institute for Molecular Imaging, University of California, Los Angeles; Department of Anesthesiology and Pain Medicine, University of Washington; The Saban Research Institute of Children’s Hospital Los Angeles; Department of Surgery, University of Southern California; Department of Biochemistry & Molecular Biology, Keck School of Medicine, University of Southern California; Division of Cardiology, Department of Medicine, University of California, Los Angeles; Molecular Biology Institute, University of California, Los Angeles; Eli and Edythe Broad Center of Regenerative Medicine and Stem Cell Research, University of California, Los Angeles; Department of Pharmacology, University of California, Los Angeles

## Abstract

The mammalian heart switches its main metabolic substrate from glucose to fatty acids shortly after birth. This metabolic switch coincides with the loss of regenerative capacity in the heart. However, it is unknown whether glucose metabolism itself regulates heart regeneration. Here, we report that glucose metabolism is a determinant of regenerative capacity in the neonatal mammalian heart. Cardiac-specific overexpression of Glut1, the embryonic form of constitutively active glucose transporter, resulted in an increase in glucose uptake and concomitant glycogen storage in postnatal heart. Upon cryoinjury, Glut1 transgenic hearts showed higher regenerative capacity with less fibrosis than non-transgenic control hearts. Interestingly, flow cytometry analysis revealed two distinct populations of ventricular cardiomyocytes: Tnnt2-high and Tnnt2-low cardiomyocytes, the latter of which showed significantly higher mitotic activity in response to high intracellular glucose in Glut1 transgenic hearts. Metabolic profiling shows that Glut1-transgenic hearts have a significant increase in the glucose metabolites upon injury, and inhibition of the nucleotide biosynthesis abrogated the regenerative advantage of high intra-cardiomyocyte glucose level. Our data suggest that the increased in glucose metabolism promotes cardiac regeneration in neonatal mouse heart.

## Introduction

Heart disease is the leading cause of death worldwide. This is in part due to the fact that the heart is the least regenerative of organs in the body. As a result, the loss of cardiomyocytes is compensated by the increase in the workload of the remaining cardiomyocytes. However, recent studies revealed that postnatal cardiomyocytes can renew at a very low but measurable rate. Neonatal mouse heart can fully regenerate up to 15% of the muscle during the first 7 days of life^1^. In human heart, cytokinesis of cardiomyocytes is detected until up to 20 years old resulting in a 3.4-fold increase in the number of cardiomyocytes since birth^2^. Pulse-chase study of atmospheric ^14^C from nuclear weapon tests suggests that human adult cardiomyocytes also undergo cytokinesis and its turnover rate is calculated to be 1% per year at age 25, declining to 0.45% by the age of 75^3, 4^. These observations can potentially be therapeutically exploited^5^. The mitotic activity of mammalian cardiomyocytes during the fetal stages is rapidly lost shortly after birth, a process known as terminal differentiation^6^. Rapid conversion from hyperplastic growth to hypertrophic enlargement at this stage coincides with the polyploidization. In rodents, the polyploidization usually accompanies karyokinesis (nuclear division) without cytokinesis (cell division). Remaining mononucleated myocytes may be responsible for postnatal proliferation of murine myocytes^1, 7, 8^. Not only the nuclear dynamics, but the response to extracellular signals also changes before and after terminal differentiation.

Interestingly, the loss of cardiac regenerative capacity coincides with the metabolic switch of energy source. The heart consumes the most energy per gram of tissue in the body, and the cardiac muscle develops a unique structure and biochemical machinery to meet its high energy demand. The embryonic heart depends mainly on glucose for ATP production, while in adult heart more than 95% of cardiac ATP is generated from fatty acid oxidation. However, during physiological embryogenesis in mice, cardiac glucose uptake drastically decreases from E10.5. This timing is long before the metabolic switch from glucose to fatty acid after birth. We have previously found that the anabolic biosynthesis of building blocks from glucose, but not the catabolic extraction of energy, is a key regulator of cardiac maturation during late embryogenesis, raising an intriguing possibility that glucose is not merely to meet the energy demand but to drive the genetic program of cardiac maturation^9^.

While recent studies have found that glucose level regulates cardiogenesis, little is known about whether and how glucose impacts the regenerative capacity of the heart *in vivo*. In this study, we report that an increase in the glucose metabolism in αMHC-hGLUT1 transgenic mice potentiates the regenerative capacity and inhibits fibrosis upon neonatal heart injury. Interestingly, we found two distinct populations of cardiomyocytes: Tnnt2^high^ and relatively immature Tnnt2^low^ cardiomyocytes. Tnnt2^low^ cardiomyocytes display a 3-4-fold increase in the mitotic activity and 2-fold increase in number in response to the increase metabolism. Glucose metabolites are significantly increased in injured Glut1 transgenic hearts, and the regenerative advantage of Glut1 transgenic hearts was abrogated by inhibition of the nucleotide biosynthesis pathway, suggesting that glucose promotes cardiac regeneration through the supply of nucleotides. Together, our results demonstrate that glucose metabolism is a determinant of regenerative capacity in neonatal mammalian heart.

## Results

### Glut1 transgenic hearts show higher glucose uptake at the basal level

Glucose is among the most tightly regulated nutrients. To increase the intra-cardiomyocyte glucose metabolism, we used *αMHC-hGLUT1* transgenic mouse line^10^. This mouse line overexpresses human GLUT1 in a cardiac-specific manner. Glut1 is an insulin-independent constitutively active glucose transporter expressed in the embryonic heart. In the postnatal heart, glucose transport is regulated predominantly by insulin-dependent Glut4. However, Glut1 becomes re-expressed in the heart upon hypertrophic stresses^11–13^. Although Glut1 is subject to heavy post-translational regulation, immunostaining indicates that GLUT1 transgene is expressed strongly in the cardiac sarcolemma in this mouse line (Supplementary Fig. 1a). Autoradiographic imaging and quantification of ^18^F-FDG-injected mouse heart suggest that ^18^F-FDG accumulation is increased by 2.6-fold in Glut1 transgenic hearts (Glut1 tg) at P2 (Supplementary Fig. 1b, c). Consistent with the original report^10^, Glut1 transgenic mice and their wild-type (WT) control litters showed no significant difference in the body weight and heart weight (Supplementary Fig. 1d and Fig. 1a). Plasma glucose level was lower but within the normal limits in transgenic mice (Supplementary Fig. 1e). Thus, *αMHC-hGLUT1* transgenic mice are suitable for analyzing the effect of hightened glucose uptake in the heart.

**Figure 1.**
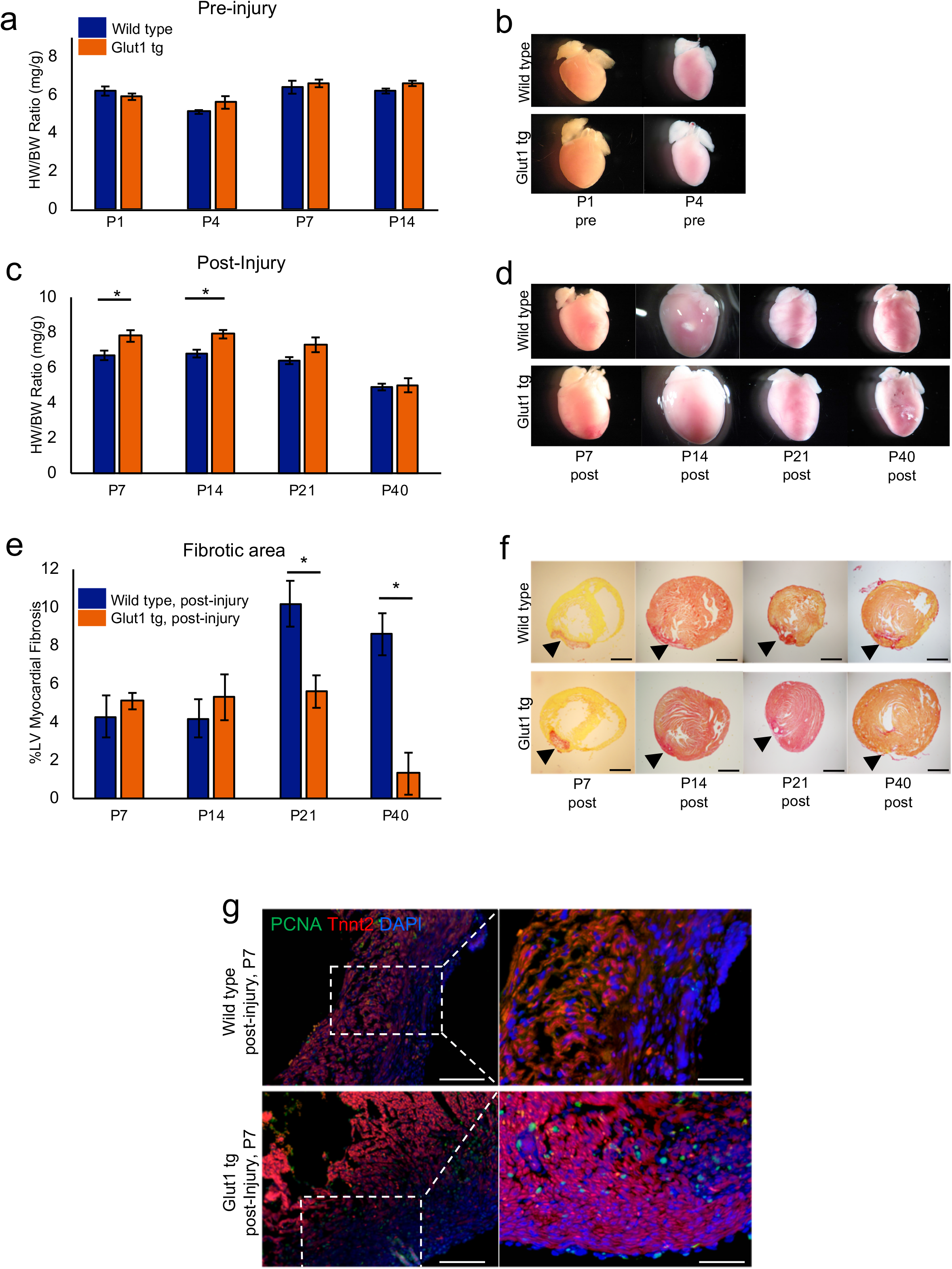
Increase in intracellular glucose promotes cardiac regeneration in Glut1 transgenic heart. a. Heart weight-body weight (HW/BW) ratio of Wild type and Glut1 transgenic (Glut1 tg) mice without injury. n = 3-7 for each group, *p* =n.s. b. Representative images of hearts. c. Heart weight-body weight (HW/BW) ratio of Wild type and Glut1 transgenic mouse post-injury. The hearts were cryoinjured at P1 and examined at P7, 14, 21 and 40. Note that the HW/BW is higher in Glut1 transgenic hearts at P7 and 14. n = 4-9 for each group, \**p* <0.05. d. Images of representative hearts post-injury. Note the ballooning of the heart at P14 in both wild type and Glut1 transgenic hearts. e. %Fibrotic area measured by Image J capture of Picrosirius red stainings of the hearts. Hearts were cryoinjured at P1 and examined at P7, 14, 21 and 40. n = 5-8 for each group, \**p* <0.05. f. Representative images of Picrosirius red staining of wild type and Glut1 transgenic hearts. Arrowheads indicate fibrotic area. Scale bar = 200 mm. g. PCNA staining of the sections from wild type and Glut1 transgenic hearts 7 days post-injury. Sections were stained with a cardiac marker (Tnnt2; Red), proliferation marker (PCNA; Green) and a nuclear marker (DAPI; Blue). Note that PCNA staining is more abundant in Glut1 transgenic heart.

### Glut1 transgenic hearts are more regenerative after injury

To test whether intra-cardiomyocyte glucose metabolism impacts the heart repair, Glut1 transgenic hearts were cryoinjured at P1 as described^14, 15^. While there was no difference in the heart weight/body weight (HW/BW) ratio between wild type (WT) and Glut1 transgenic hearts without injury (Fig. 1a, b), Glut1 transgenic hearts showed a statistically significant increase in the HW/BW 7 and 14 days post-injury (Fig. 1c, d; 7.8 ± 0.33 vs 7.9 ± 0.24, p < 0.05). Glut1 transgenic hearts continued to show a higher HW/BW at day 21 but no statistically significant differences were seen by day 40 post-injury. To examine the collagen deposition post-injury, the hearts were sectioned and analyzed by H&E and Picrosirius Red stainings. Image-J quantification of scar area/left ventricular area showed 4-5% fibrotic area in both WT and Glut1 transgenic hearts at day 7 and 14 post-injury. Cryoinjury induced up to 8-10% fibrotic area in WT hearts at P21 and P40, whereas Glut1 transgenic hearts showed a significantly lower level of fibrosis (Fig. 1e, f; 10.2±1.2 vs 5.6 ± 0.85 p < 0.01, at P21). To determine whether the decreased level of fibrosis was associated with proliferation of cardiomyocytes and thus a beneficial effect of high intra-cardiomyocyte glucose, the WT and Glut1 transgenic hearts were immunostained for PCNA (mitosis marker) and Tnnt2 (cardiac marker). Glut1 transgenic hearts showed an increase in the number of PCNA-positive cardiomyocytes (Fig. 1g). Cardiomyocyte proliferation and angiogenesis are both required for complete regeneration of the neonatal heart after injury^16^. In our study, we found that neovascularization was increased around the border zone of the Glut1 transgenic hearts when compared to the WT control hearts (Supplementary Fig. 3). The increase in vascularity is likely secondary to the increase in cardiomyocyte proliferation, as Glut1 is not overexpressed in the vasculature in our experiments. Echocardiographic analysis revealed no significant difference in the cardiac function between WT and Glut1 transgenic heart at P21 and 40 (Supplementary Fig. 2). Together, these data suggest that Glut1 transgenic hearts show higher regenerative capacity possibly due to the increased mitotic activity of cardiomyocytes.

**Figure 2.**
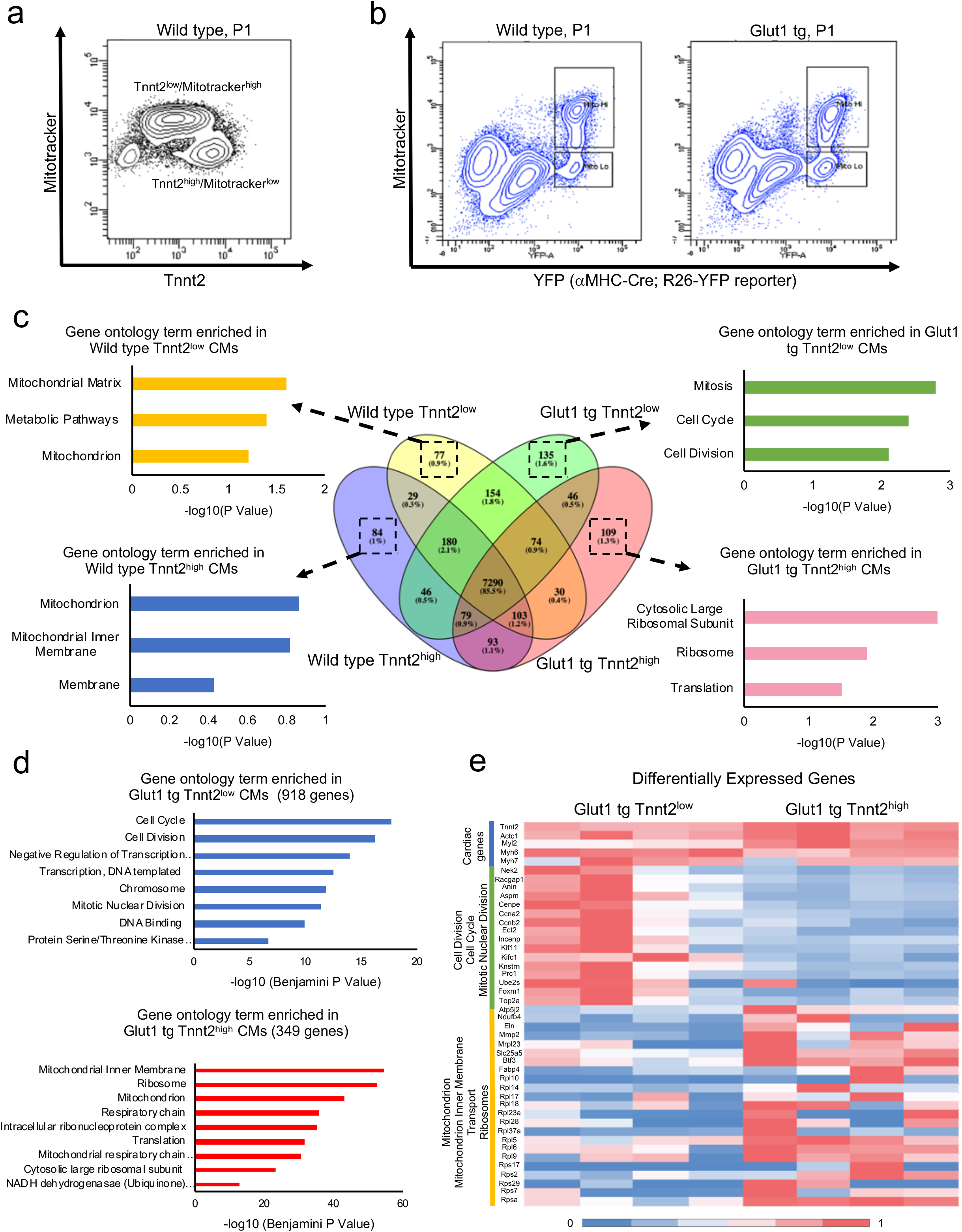
mRNA expression profile of Tnnt2^high^ and Tnnt2^low^ cardiomyocytes from Wild type and Glut1 transgenic hearts at P1. a. Representative dot plot of flow cytometry analysis of P1 Wild type hearts for Tnnt2 (cardiac marker) and Mitotracker (mitochondrial contents). Note two populations of cardiomyocytes. b. Flow cytometry analysis of P1 *aMHC-Cre* ^*tg*^; *R26^+/YFP reporter^* and *aMHC-hGLUT1* ^*tg*^; *aMHC-Cre* ^*tg*^; *R26^+/YFP reporter^* analysis for YFP (cardiac lineage) and Mitotracker (mitochondrial contents). Note two populations of cardiomyocytes. c. Venn diagram of mRNA expressed in Tnnt2^high^ and Tnnt2^low^ cardiomyocytes from Wild type and Glut1 transgenic hearts. Bar graphs represent top three gene ontology (GO) term enriched in the genes uniquely expressed in each population. Although 4 populations are similar, G1 Tnnt2^low^ cardiomyocytes are enriched for the genes associated with mitosis, cell cycle, and cell division. d. GO analysis of the genes upregulated in Glut1 tg Tnnt2^low^ vs Glut1 tg Tnnt2^high^. e. Heatmap of expression level of representative cardiac, cell cycle, and mitochondrial genes differentially expressed in Glut1 tg Tnnt2^low^ vs Glut1 tg Tnnt2^high^.

### Two distinct cardiomyocytes populations in neonatal hearts

As higher mitotic activity of fetal/neonatal cardiomyocytes often accompanies lower maturation level during cardiogenesis^9^, we characterized the maturity of cardiomyocytes by Tnnt2 expression (a cardiac specific contractile protein) and Mitotracker (a marker of mitochondrial content). Flow cytometric analysis revealed two distinct populations of cardiomyocytes in neonatal hearts; Tnnt2^high^ and Tnnt2^low^ cardiomyocytes (Fig. 2a). Tnnt2^high^ and Tnnt2^low^ cardiomyocytes were Mitotracker^low^ and Mitotracker^high^, respectively. As the Mitotracker does not necessarily reflect the mitochondrial function^17–19^, we examined the mRNA expression signature of Tnnt2^high^ and Tnnt2^low^ cardiomyocytes genome-wide. To collect the cells alive, two populations were sorted from *αMHC-Cre* ^*tg*^; *R26*^+/*YFP reporter*^ hearts in both WT and Glut1 transgenic background using YFP reporter and Mitotracker (Fig. 2b). Forward scatter plot suggests that Tnnt2^low^ (Mitotracker^high^) cardiomyocytes are slightly smaller in size (Supplementary Fig. 4c). RNA-seq analysis confirmed that Tnnt2^low^ (Mitotracker^high^) cardiomyocytes do express a slightly lower level of Tnnt2 mRNA as compared to Tnnt2^high^ (Mitotracker^low^) cardiomyocytes. Although Mitotracker staining level is high, mitochondrial gene expression level was lower in Tnnt2^low^ (Mitotracker^high^) cardiomyocytes than Tnnt2^high^ (Mitotracker^low^) cardiomyocytes in WT heart, suggesting that Tnnt2^low^ (Mitotracker^high^) cardiomyocytes represent relatively immature cardiomyocytes and that Mitotracker does not reflect the functional maturity of mitochondria in this context, consistent with previous observations^17–19^.

To examine how the increase in intra-cardiomyocyte glucose uptake impacts the expression signature of Tnnt2^high^ and Tnnt2^low^ cardiomyocytes, the RNA-seq was also performed in these populations in Glut1 transgenic hearts. Comparison of 4 populations (Tnnt2^high^ and Tnnt2^low^ cardiomyocytes from WT and Glut1 hearts) showed only marginal differences at transcriptome level (Fig. 2b). Interestingly, however, pathway analysis revealed that Tnnt2^low^ cardiomyocytes from Glut1 transgenic heart (G1 Tnnt2^low^) are unique in that they are enriched for the genes associated with mitosis, cell cycle, and cell division compared to the other 3 populations in WT and Glut1 transgenic hearts (Fig. 2b). Indeed, 2-way comparison between G1 Tnnt2^low^ and G1 Tnnt2^high^ cardiomyocytes identified considerably more differentially expressed genes than any 2 comparisons (918 and 349 genes up/down-regulated in G1 Tnnt2^low^ cells; Supplementary Fig. 4a). Compared to G1 Tnnt2^high^ cardiomyocytes, G1 Tnnt2^low^ cardiomyocytes are, again, enriched for the genes associated with cell cycle, cell division, and mitotic nuclear division, while cardiac genes and mitochondrial genes are expressed at a lower level (Fig. 2d, e). Together, we found two distinct populations of cardiomyocytes in neonatal heart: Tnnt2^high^ cardiomyocytes are larger, less mitotic and more mature than Tnnt2^low^ cells. On the other hand, Tnnt2^low^ cells show less mature expression signature and upregulate cell cycle gene expression in Glut1 transgenic background.

### Tnnt2-low cardiomyocytes are glucose-responders

As the cell cycle genes are enriched in WT Tnnt2^low^ cardiomyocytes and further upregulated upon the increase in glucose uptake, the mitotic activity of Tnnt2^high^ and Tnnt2^low^ cardiomyocytes were tested by EdU incorporation assay. The heart was injured at P1, and EdU was injected intraperitoneally at P7 followed by cardiomyocyte isolation and flow cytometry analysis for Tnnt2, Mitotracker, and EdU signals. Consistent with the gene expression profile (Fig. 2), mitotic index (%EdU incorporation) was significantly higher in G1 Tnnt2^low^ cardiomyocytes than WT Tnnt2^low^, whereas G1 Tnnt2^high^ cardiomyocytes were not significantly more mitotic than WT Tnnt2^high^ cardiomyocytes (Fig. 3a-c). Thus, consistent with the transcriptome data (Fig. 2), Tnnt2^low^ subset of neonatal cardiomyocytes are glucose-responders that are mitotically potentiated in response to the increase in glucose uptake in the Glut1 transgenic background.

**Figure 3.**
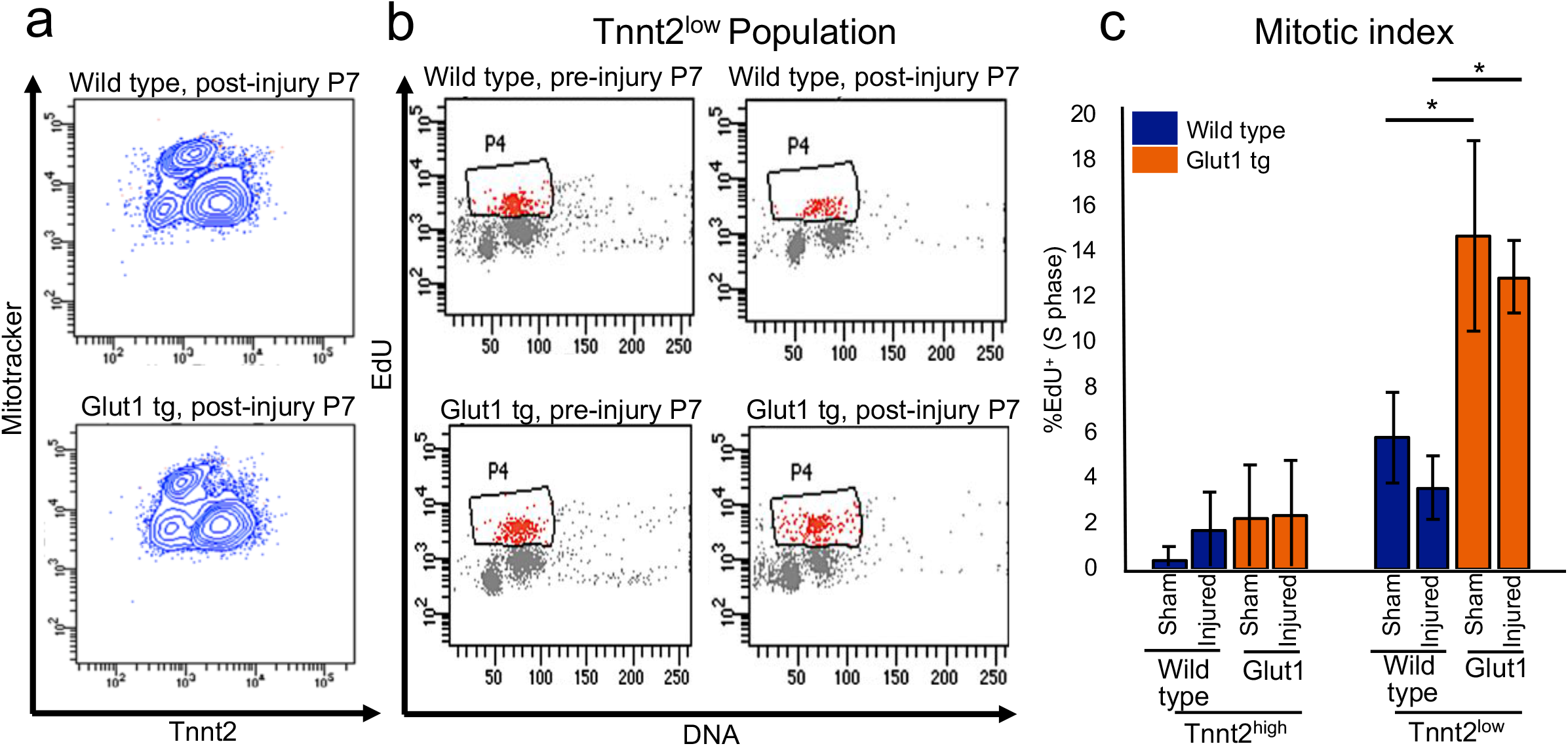
Tnnt2^low^ cardiomyocytes are mitotically activated by the increase in intracellular glucose. a. Representative flow cytometry profile of the ventricular cardiomyocytes from P7 Wild type and Glut1 transgenic hearts stained with Tnnt2 and Mitotracker. b. Representative flow cytometry plot of EdU incorporation assay of Tnnt2^low^ cardiomyocytes. Wild type and Glut1 transgenic mice were injected with EdU and the ventricular cardiomyocytes were isolated and analyzed by Tnnt2, Mitotracker, and EdU staining. c. Mitotic index of Tnnt2^high^ and Tnnt2^low^ cardiomyocytes in Wild type and Glut1 transgenic heart with or without injury. n = 3-8. \**p*<0.05. \*\*\**p*<0.001. Note that Tnn2^low^ cardiomyocytes are more mitotic than Tnn2^high^ cardiomyocytes and that the mitotic activity of Tnnt2^low^ cardiomyocytes drastically increases in Glut1 transgenic background.

### Metabolomics analysis suggests an increase in glucose metabolites upon injury in Glut1 transgenic heart

Glucose is metabolized in multiple pathways including glycolysis, hexosamine biosynthesis pathway, and the pentose phosphate pathway (PPP). Having demonstrated that the increase in cardiac glucose uptake potentiates the regeneration in neonatal heart, we sought for its downstream mechanism. Metabolomics analysis of uninjured Glut1 transgenic hearts showed a significant increase in nucleotides levels including AMP and CMP (Fig. 4a, Supplementary Fig. 5). Although the intracellular glucose level is not increased, Periodic Acid Schiff (PAS) staining showed significant accumulation of glycogen in Glut1 transgenic hearts, indicating that excess glucose is partly stored in the form of glycogen (Fig. 4b). Upon injury, metabolites in the PPP and glycolysis were increased in Glut1 transgenic hearts. In particular, metabolites in polyol pathway (sorbitol/mannitol, fructose) were significantly increased, indicating the active utilization of glucose upon injury.

**Figure 4.**
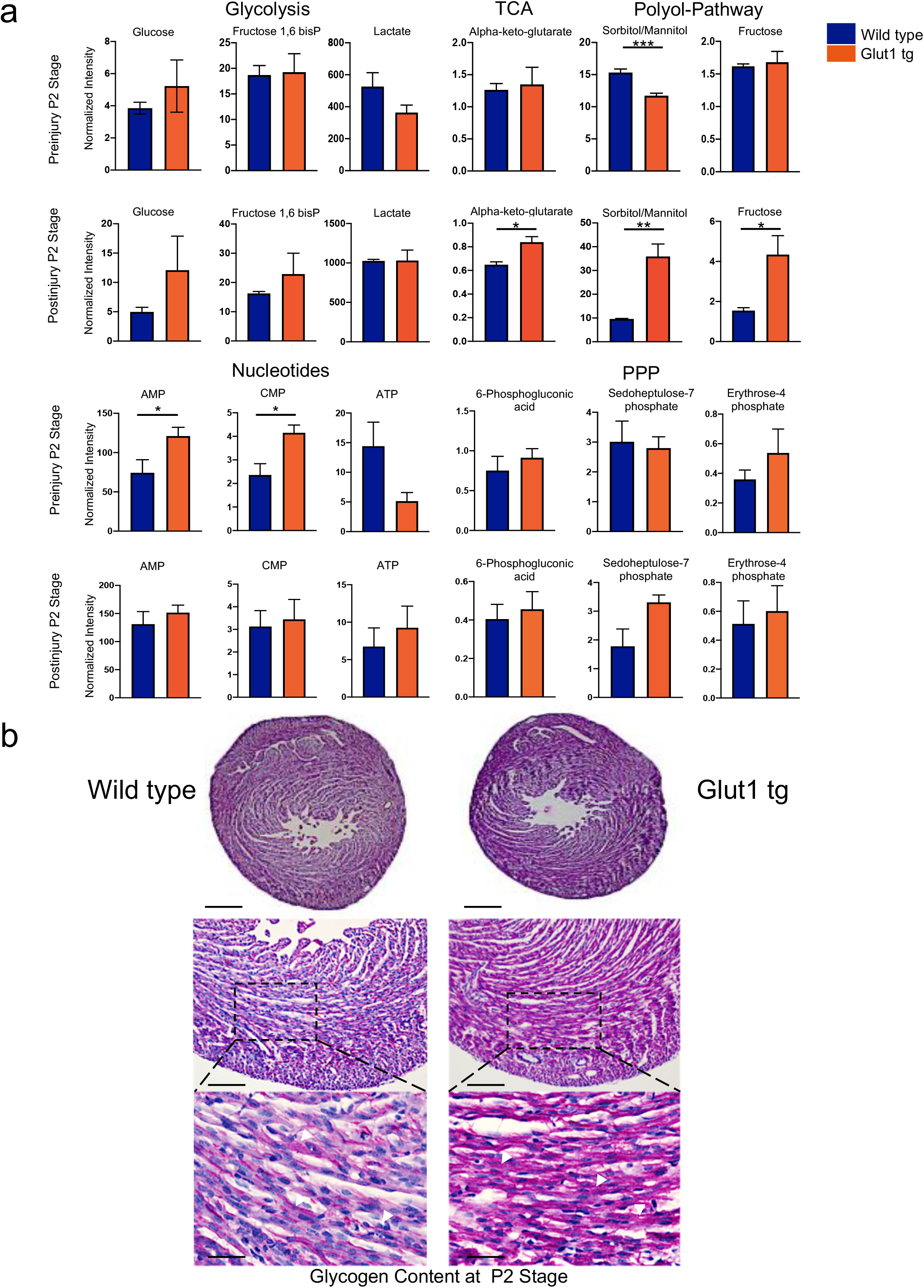
Metabolomics analysis of Glut1 transgenic hearts. a. Metabolomics analysis of Glut1 transgenic hearts and its controls from wild type litters at P2 with or without injury. Overall, the metabolic profile shows no drastic difference at the basal level, but glucose utility increases upon injury as represented by the increase in the metabolites in polyol pathway and glucose. b. Histological analysis of glycogen storage of Glut1 transgenic hearts and its controls from wild type litters at P2 by Periodic Acid Staining. Representative images of 3 samples in each group. Note the increased intracellular glycogen storage in Glut1 transgenic hearts.

**Figure 5.**
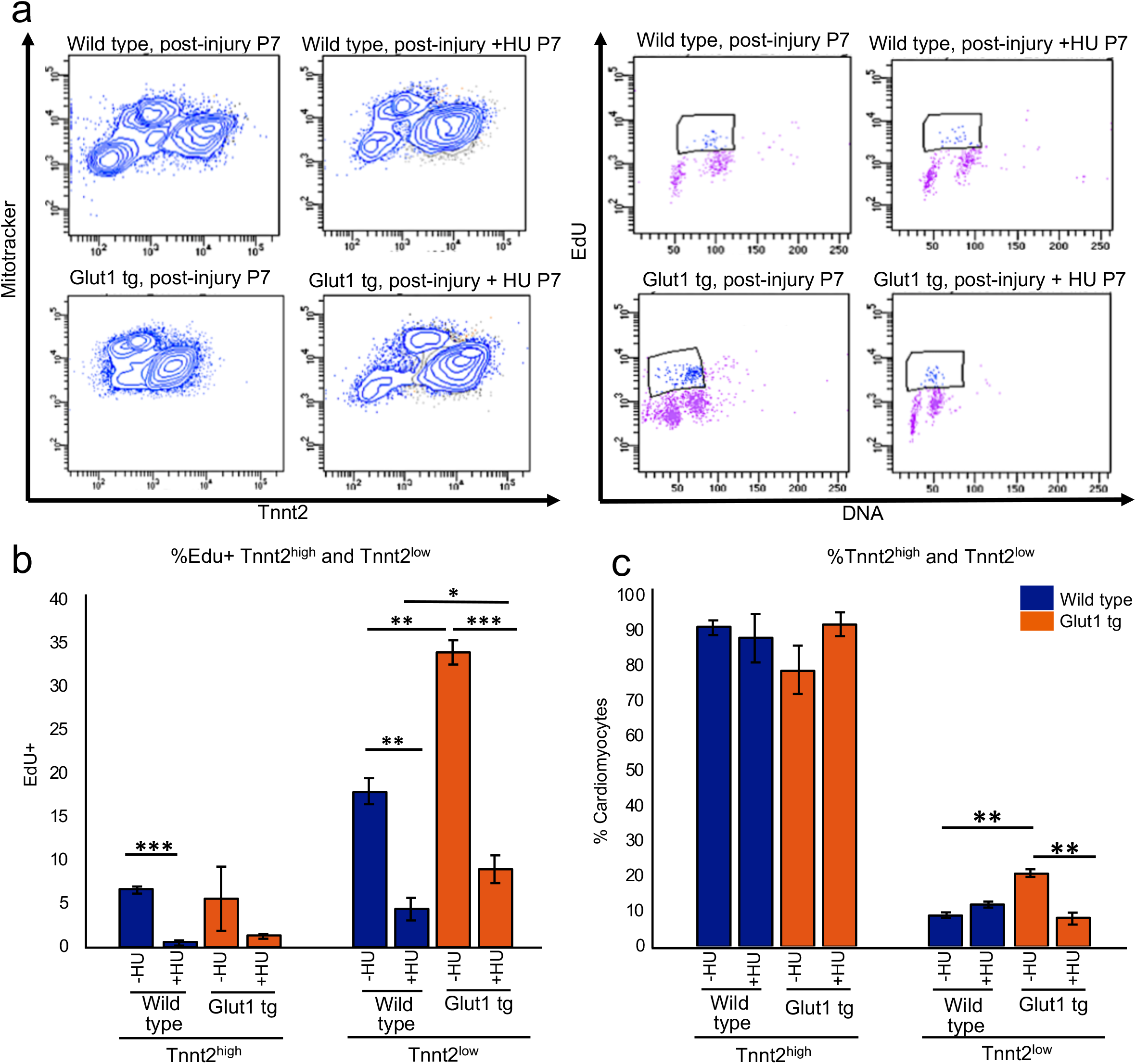
Impact of hydroxyurea on the proliferation of Tnnt2^low^ cardiomyocytes. a. Representative contour plots (left) for Tnnt2 and Mitotracker and dot plots (right) for EdU and DNA content by flow cytometry analysis at P7. Wild type and Glut1 transgenic hearts were cryoinjured at P1 and treated with daily injection of HU or vehicle until the analysis at P7. b. Quantification of %EdU positive cells in Tnnt2^high^ and Tnnt2^low^ cardiomyocytes in wild type and Glut1 transgenic hearts with or without HU treatment. Note that both Tnnt2^high^ and Tnnt2^low^ cardiomyocytes respond to HU treatment but that EdU incorporation of Tnnt2^low^ cardiomyocytes was drastically decreased by HU treatment. n = 4. \*\*\**p*<0.01. c. Quantification of the Tnnt2^high^ and Tnnt2^low^ fractions of cardiomyocytes in wild type and Glut1 transgenic hearts with or without HU treatment. While Tnnt2^high^ or Tnnt2^low^ fractions was not influenced by HU in wild type heart, HU treatment significantly decreased the percentage of Tnnt2^low^ cardiomyocytes in Glut1 transgenic hearts. n = 4. \*\**p*<0.01. \*\*\**p*<0.001.

### Inhibition of the nucleotide biosynthesis abrogates the regenerative advantage of Glut1 transgenic heart

Our previous report found that glucose inhibits the cardiac maturation and promotes proliferation via the nucleotide biosynthesis during fetal development^9^. To test whether nucleotide biosynthesis also plays a central role in the glucose-induced cardiomyocyte regeneration, the neonatal hearts were injured in the presence of hydroxyurea (HU). HU is an inhibitor of ribonucleotide reductase (RNR), the rate-limiting enzyme of the nucleotide biosynthesis, that passes through the placenta when intraperitoneally injected^20^. 10 mg/kg of HU was intraperitoneally injected daily for 7 days starting 4 hours prior to the cryoinjury at P1 until the EdU incorporation assay at P7 (Fig. 5a). HU blocked EdU incorporation in the in both Tnnt2^low^ and Tnnt2^high^ cardiomyocytes (Fig. 5b), and the number of Tnnt2^low^ cardiomyocytes was significantly reduced compared to the vehicle controls (Fig. 5c). Thus, Tnnt2^low^ cardiomyocytes contribute to cardiac regeneration in a glucose-dependent manner via the supply of nucleotides.

## Discussion

In this study, we demonstrated that glucose metabolism is a determinant of regenerative capacity of the heart. By increasing basal intra-cardiomyocyte glucose uptake using transgenic mice that specifically overexpresses Glut1, we successfully enhanced the regenerative capacity of neonatal mouse hearts. We identified two distinct subpopulations of cardiomyocytes that have not been previously described; relatively mature (Tnnt2^high^) and immature (Tnnt2^low^) cardiomyocytes. Interestingly, the mitotic activity was higher in the Tnnt2^low^ immature cardiomyocytes in transgenic Glut1 hearts. Thus, Tnnt2^low^ subset of cardiomyocytes are the glucose-responders that proliferate in response to high glucose uptake upon heart injury, resulting in acceleration of heart regeneration.

Shortly after birth, the cardiomyocytes undergo terminal differentiation, and the response to the stress and extracellular signals changes. For example, Angiotensin II (Ang-II) or phenylephrine (PE) directly stimulates hypertrophy of the heart via the ERK/MAPK pathway at postnatal stages, while the same signals trigger hyperplasia during late gestational stages^21^. Our data suggest increased glucose metabolism potentiates the hyperplastic growth upon cardiac injury at neonatal stages in Glut1 transgenic heart. In adult heart, Glut1-overexpressing cardiomyocytes seem to respond to the pressure overload predominantly by hypertrophic growth^10^. However, it is unknown whether cardiomyocyte proliferation is simultaneously stimulated in the adult Glut1 transgenic hearts. Further investigation will determine whether glucose loading strategy may promote the regeneration of adult heart.

There is generally an inverse correlation between cell proliferation and differentiation during developmental stages^22^. In our previous study, we found that the nucleotide biosynthesis supported by the PPP is a key mechanism balancing cell proliferation and maturation during the physiological growth of cardiomyocytes. The balance between proliferation and maturation was successfully manipulated by addition of glucose or nucleotides (gain-of-function) and inhibition of the PPP and nucleotide biosynthesis (loss-of-function) in the *in vitro* settings^9^. The current study demonstrates that similar intervention may be effective during the neonatal stages to accelerate regeneration. Metabolic profiling of neonate hearts revealed that glucose is actively utilized in the context of cryoinjury, evidenced by the marked accumulation of polyol pathway intermediates. The influence of metabolic activity, driven by glucose utilization, on regulating cardiac regeneration was probed further through pharmacological intervention. The regenerative advantage of Glut1 transgenic heart was abrogated by the inhibition of RNR, the key enzyme of nucleotide biosynthesis pathway, by HU, further re-emphasizing the importance of the nucleotides in regulating cardiomyocyte proliferation and regeneration upon injury.

Cells display distinct metabolic profiles depending on their differentiation stage^23–25^. The fuel type of cells serves not merely as a source of energy but also as a regulator of self-renewal and differentiation of stem or progenitor cells^26–29^. Among such metabolic factors, environmental glucose is one of the fundamental nutrients for the cells and is a significant contributor to heart formation. However, despite the established association between hyperglycemia and the heart malformation, little is known about how glucose impacts cardiomyocyte formation and re-formation^30–32^. We have recently found that glucose induces cardiomyocyte proliferation at the sacrifice of its maturity via the pentose phosphate pathway and nucleotide biosynthesis during physiological heart formation^9^. The current study is the first to demonstrate that glucose metabolism is a critical determinant of cardiac regenerative capacity *in vivo*. The non-genetic factors and genetic regulators mutually reinforce each other to establish a robust cardiac differentiation program. Metabolic environment of the cells is, on one hand, determined by the genetic program of cells. On the other hand, the genetic program of the cells is dependent on the metabolic environment. Understanding how the genetic program and metabolic factors crosstalk during physiological and pathological cardiogenesis will be an important theme that requires further investigation.

## Materials and Methods

### Mouse maintenance and cryoinjury of neonatal mouse hearts

Animals were housed at the University of California Los Angeles (UCLA) animal facility. WT and transgenic Glut1 transgenic mice were maintained on the ICR/CD-1 strain background according to the Guide for the Care and Use of Laboratory Animals published by the US National Institute of Health (NIH Publication No. 85-23, revised 1996). Housing and experiments were performed according to the Institutional Approval for Approval for Appropriate Care and Use of Laboratory Animals by the UCLA Institutional Animal Care and Use Committee (Protocol # 2008-143-31). Some WT ICR/CD-1 mom and pups were ordered from Charles River Laboratories, MA and housed at the UCLA animal facility. Surgical procedures were performed as previously described^33^ with some modifications in order to optimize the time of injury to the apex of the left ventricle while creating a non-transmural injury as well as optimizing the time for adequate anesthesia and decreasing post-surgical mortality of the pups. Briefly, 1-day old neonatal mice were anesthetized under hypothermic conditions. Left lateral thoracotomies at the 5^th^ intercostal space were performed and hearts were exposed through an incision at intercostal space 5-6. As soon as the incision was made, a 1mm metal probe was chilled in liquid nitrogen for 15-30 seconds. After heart exposure, the pre-chilled probe was placed on the apex of the left ventricle for 1 second. Sham operated hearts were also exposed for 1 second but no injury was done. The thoracic wall incision was closed with a 7-0 Prolene suture (8701H, Ethicon, Somerville, NJ USA) using a figure of eight suture technique. The skin was closed by topical tissue adhesive (Gluture, Abbott Laboratories, IL, USA). After surgery, pups were placed immediately in a warm water bed and allowed to full recovery for at least 15 minutes prior to putting pups back in the cage with mom. The survival rate was around 90%. Body weights were measured (grams) using a weight scale prior to each cryosurgery and plasma glucose levels were measured by using a ONE TOUCH Glucose Monitor using blood samples obtained by tail bleeding.

### Tissue collection and histological analysis

To collect hearts at different time points, mice were put under general anesthesia with 99.9% isoflurane in an inhalation chamber followed by cervical dislocation. Body weights were measured prior to collection of hearts. After collection of hearts, they were weighed using an analytical balance weight scale. Hearts were dissected and cleaned prior to weighting. Pictures were taken for each heart in order to analyze phenotypic differences between the transgenic Glut1 hearts and the WT hearts. Hearts were embedded in paraffin and sectioned transversely from the apex to the base at 7μm. Paraffin sections from hearts at 1, 2, 3 and 6 weeks after surgery were stained with Hematoxylin and Eosin according to standard protocols to evaluate the extent of injury. The heart sections were also stained with Picrosirius Red Staining to order to determine the extent of collagen deposition. After deparaffinization in Xylene, the sections were washed with ethanol and PBS, incubated with picrosirius red solution for 1 hour at room temperature, and then rinsed quickly with acetic acid and ethanol. Stained sections were dehydrated with ethanol and xylene and mounted. In Picrosirius Red stained sections, the scar tissue is stained red and muscle tissue appears yellow. For immunostaining, sections were incubated with the following primary antibodies: cardiac TnnT (Sigma Cat# HPA015774 1:1000 rabbit), Proliferating Cell Nuclear Antigen (Vector laboratories PCNA Cat#VP-P980), and PECAM(eBioscience Cat# 13-0311) for 1 hour. Heart sections were also stained with Periodic Acid Schiff stain kit (Abcam) at post-natal stage 2 days in order to determine the glycogen content within these hearts. Glycogen stained magenta in color while nuclei stained blue.

### Flow cytometry

Hydroxyurea (HU) and Edu were injected intraperitoneally. HU injections were performed 4 hours prior to cryoinjury and daily for 7 days. Edu was injected 4 hours prior to tissue harvest, hearts were dissociated to single cells at P7 as previously described^9, 34^. The cells were stained for MitoTracker Orange CMTMRos (ThermoFisher), and further fixed and strained with Tnnt2 antibody (rabbit, 1:200, Sigma-Aldrich) and ClickIT EdU kit. Alexa488-conjugated anti-rabbit IgG antibody (ThermoFisher) was used for secondary antibody for TnnT2. Stained cells were analyzed by a flow cytometer (LSRII, BD Biosciences). Data analysis was performed using FACSDiva (BD Biosciences).

### RNA-Seq and Data Analyses

Tnnt2^high^ and Tnnt2^low^ cardiomyocytes from *αMHC-Cre^tg^; R26^+/YFP reporter^* and *αMHC-hGLUT1^tg^; αMHC-Cre^tg^; R26^+/YFP reporter^* were sorted using Mitotracker and YFP and their RNAs were extracted using RNeasy kit (QIAGEN). mRNA library was prepared with the Illumina HiSeq mRNA kit (Illumina, RS-122-2001), according to manufacturer’s instructions. Final libraries were sequenced with single-end reads of 50bp on Illumina sequencing platforms HiSeq 3000 (Illumina, San Diego, CA); raw sequencing reads were assessed for quality using FastQC (www.bioinformatics.babraham.ac.uk/projects/fastqc), then low-quality bases with a Phred quality value lower than 20 were trimmed off the read ends. The RNA Sequencing reads were mapped to mouse genome assembly UCSC mm10 using the HISAT2 with default settings^35^. UCSC annotation release mm10 was used to identify genes for feature counting. Read counts for genes were determined as counts mapped to the union of all exons for all transcript isoforms of a given gene using HTSeq^36^. Differential expression analysis and generation of the corresponding data plots were performed using DESeq2 using default settings, except that the alpha (FDR) threshold was set to 0.05^37^.

### 18 F-FDG measurement by Autoradiography

Autoradiography was conducted as we described^38^ except that mouse pups were injected with 18F-FDG (3.33 MBq; UCLA Ahmanson Biomedical Cyclotron Facility) and hearts were removed and sliced at 7um of thickness for analysis. Heart ^18^F-FDG accumulation was quantified using Fujifilm Multi Gauge v.3.0 software and the signal was normalized to the amount of probe injected for each mouse pup.

### Intracellular metabolite extraction and analysis

Neonatal hearts were excised and weighed. Hearts were submerged in 500 μl of extraction solution (80% methanol / 10uM trifluoromethanosulfanate) contained in 2 mL tubes containing 1.4 mm ceramic beads. Tubes were placed in a Bead Mill 4 (Fisherbrand) to homogenize tissue for 6 minutes (speed: 6 m/s). Homogenates were transferred to clean tubes and centrifuged at 4 °C for 10 min at 17,000g. For every sample, a supernatant volume that is equivalent to 5 mg of total heart tissue was transferred to a glass vial. Samples in glass vials were dried under an EZ-2Elite evaporator. Metabolites were resuspended in 100 μl 50% acetonitrile (ACN). The analysis was performed on a Q Exactive (Thermo Scientific) in polarity-switching mode with positive voltage 3.5 kV and negative voltage 3.5 kV. The mass spectrometer was coupled to an Vanquish (Thermo Scientific) UHPLC system. Mobile phase A was 5 mM NH_4_Ac, pH 9.9, B was ACN. Separation was achieved on a Luna 3 mm NH2 100 A (150 × 2.0 mm) (Phenomenex) column kept at a temperature of 40 °C. Injection volume was 10 μl, flow was 200 μl min^−1^, and the gradient ran from 15% A to 95% A in 18 min, followed by an isocratic step for 9 min and re-equilibration for 7 min. Metabolites were detected and quantified as the area under the curve based on retention time and accurate mass (≤15 p.p.m.) using MZMine 2.0.

### Quantification and Statistics

Data are presented as mean +/− SEM. Comparison between the two groups was done by the two tailed Student’s T test. *P<0.05 and **P<0.01 were considered statistically significant. Quantification of scar size as a ratio of scar area to the left ventricular area was conducted manually by using Image J software by one observer, however the observer was not blinded to the experimental groups.

## Acknowledgement

Authors thank technical support from Dr. Morizono (AIDS institute, UCLA) and BSCRC FACS core. Authors are also grateful to Dr. Sherin Devaskar, Dr. Marlin Touma, and Dr. Josephine Enciso for helpful comments. This study was supported by NIH HL142801 to AN, UCLA David Geffen School of Medicine (DGSOM) Seed Grant to AN and HC, NIH HL130172 to CLL, UCLA Eli and Edythe Broad Center of Regenerative Medicine and Stem Cell Research (BSCRC) Training Grant and Children’s Discovery and Innovation (CDI) Grant to VMFM.

---

**Supplementary Figure 1.**
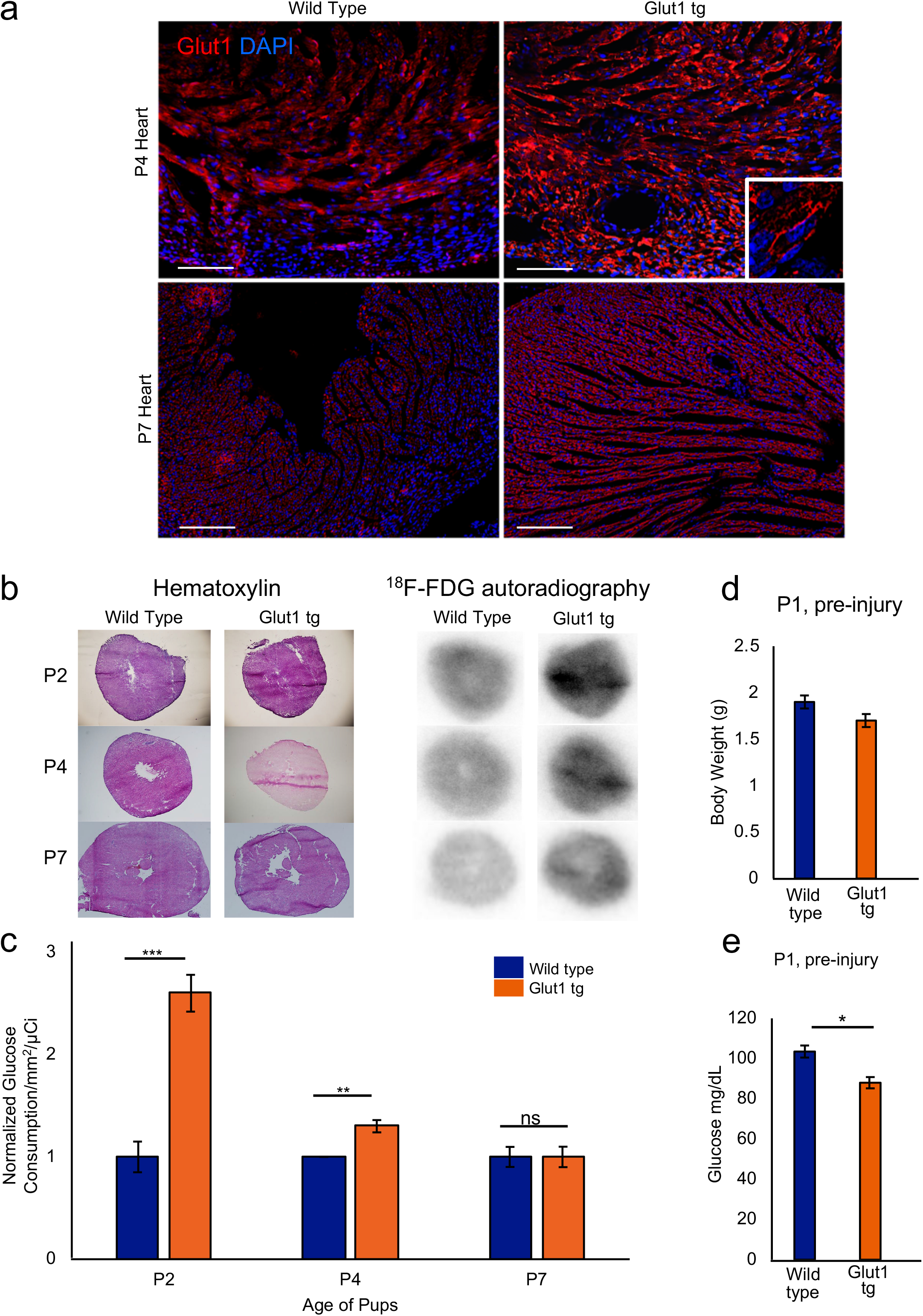
Increased glucose uptake in Glut1 transgenic heart. a. Glut1 immunofluorescent staining of *aMHC-hGLUT1* transgenic heart at P4 and P7. Glut1 transgene is expressed in the sarcolemma. b. Glucose uptake of Glut1 transgenic hearts measured by autoradiography of accumulated ^18^F-FDG activity at P2, 4, and 7. c. Glucose uptake of Glut1 transgenic heart measured by ^18^F-FDG activity in the autoradiography images at P2, 4, and 7. n = 1-3 for each group** p<0.01, *** p<0.001 d. Body weight of at P1, pre-injury, n = 28 vs 29, *p*=n.s. by t-test. e. Blood glucose at P1, pre-injury, n = 28 vs 29, \**p*<0.05 by t-test.

**Supplemental Figure 2.**
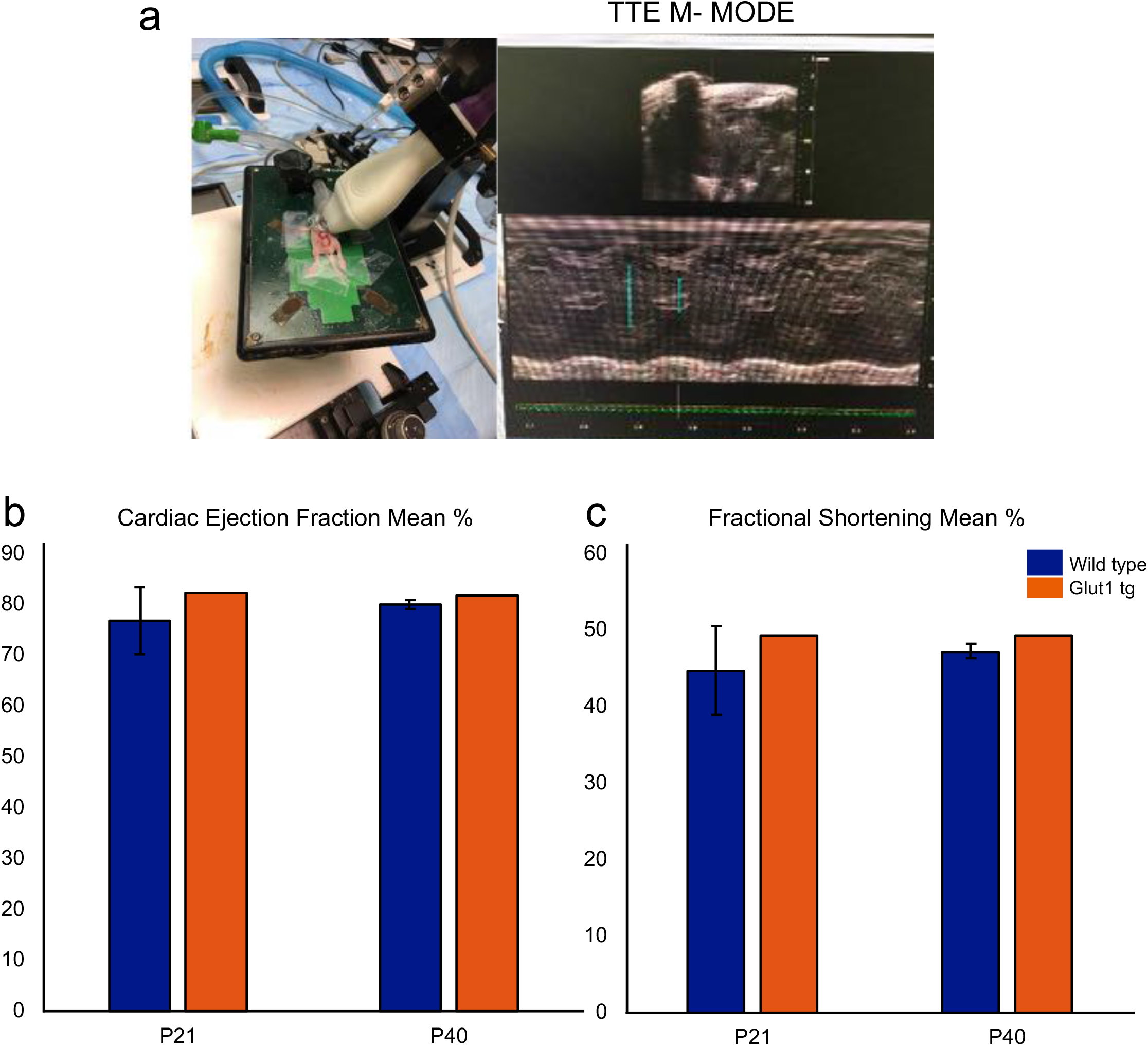
Cardiac function of Wild type and Glut1 transgenic hearts after surgery. a. In vivo trans-thoracic echocardiography and representative M mode echocardiogram. b. Cardiac ejection fraction at P21 and P40 stages after surgery. P21 stage n = 3, each. P40 stage n = 3 wild type, n = 1 Glut1 tg c. Fractional shortening at P21 and P40 stages after surgery. P21 stage n = 3, each. P40 stage n = 3 wild type, n = 1 Glut1 tg All data represent mean +/− SEM.

**Supplementary Figure 3.**
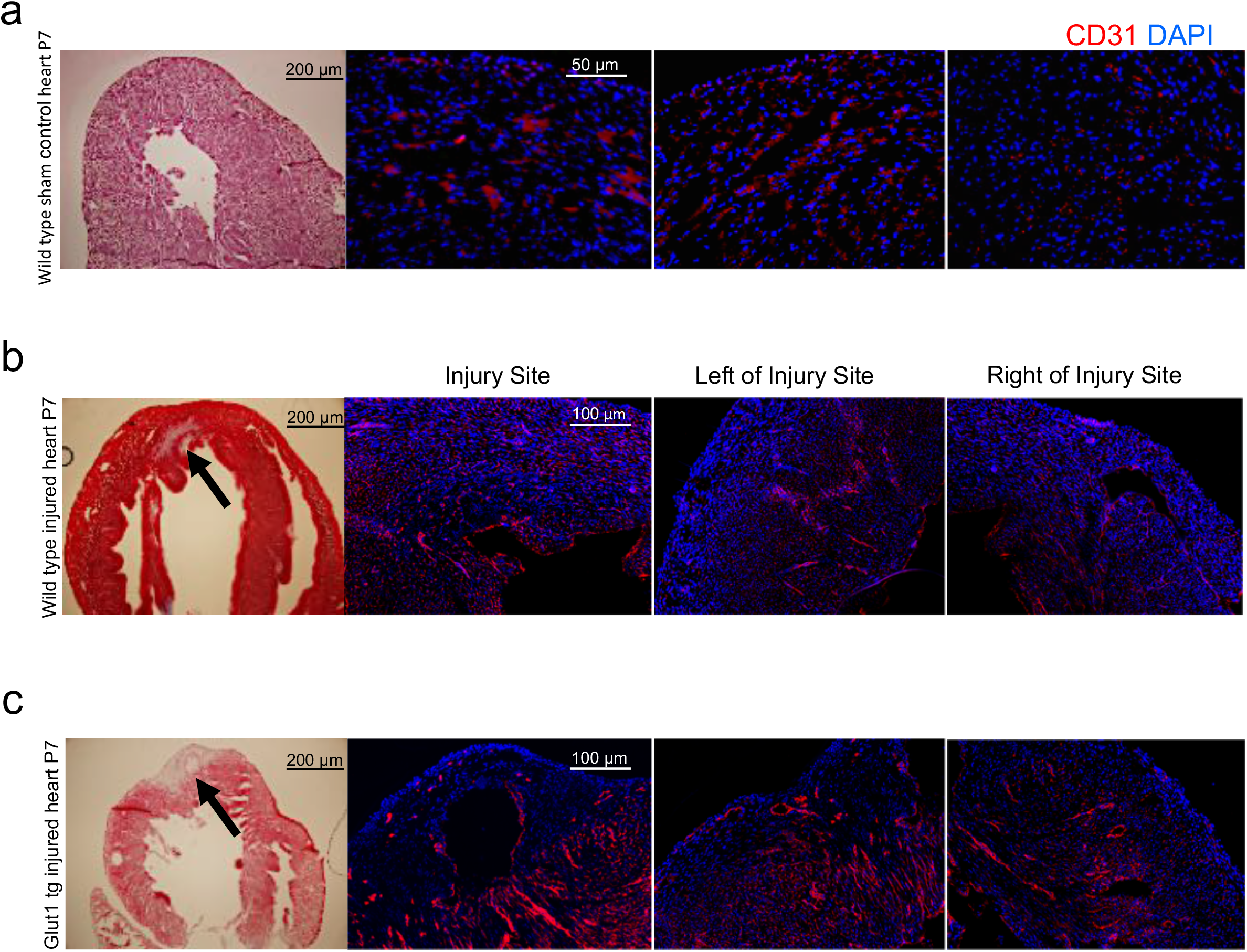
Increased neovascularization in the border zone of Glut1 hearts after cardiac injury. Representative images of Wild type sham (a), Wild type injured (b), and Glut1 transgenic injured (c) hearts at p7. Left, (a) representative image of H&E staining, (b,c) representative image of Masson’s Trichome staining. Right, immunostaining with CD31 (endothelial marker; red).

**Supplementary Figure 4.**
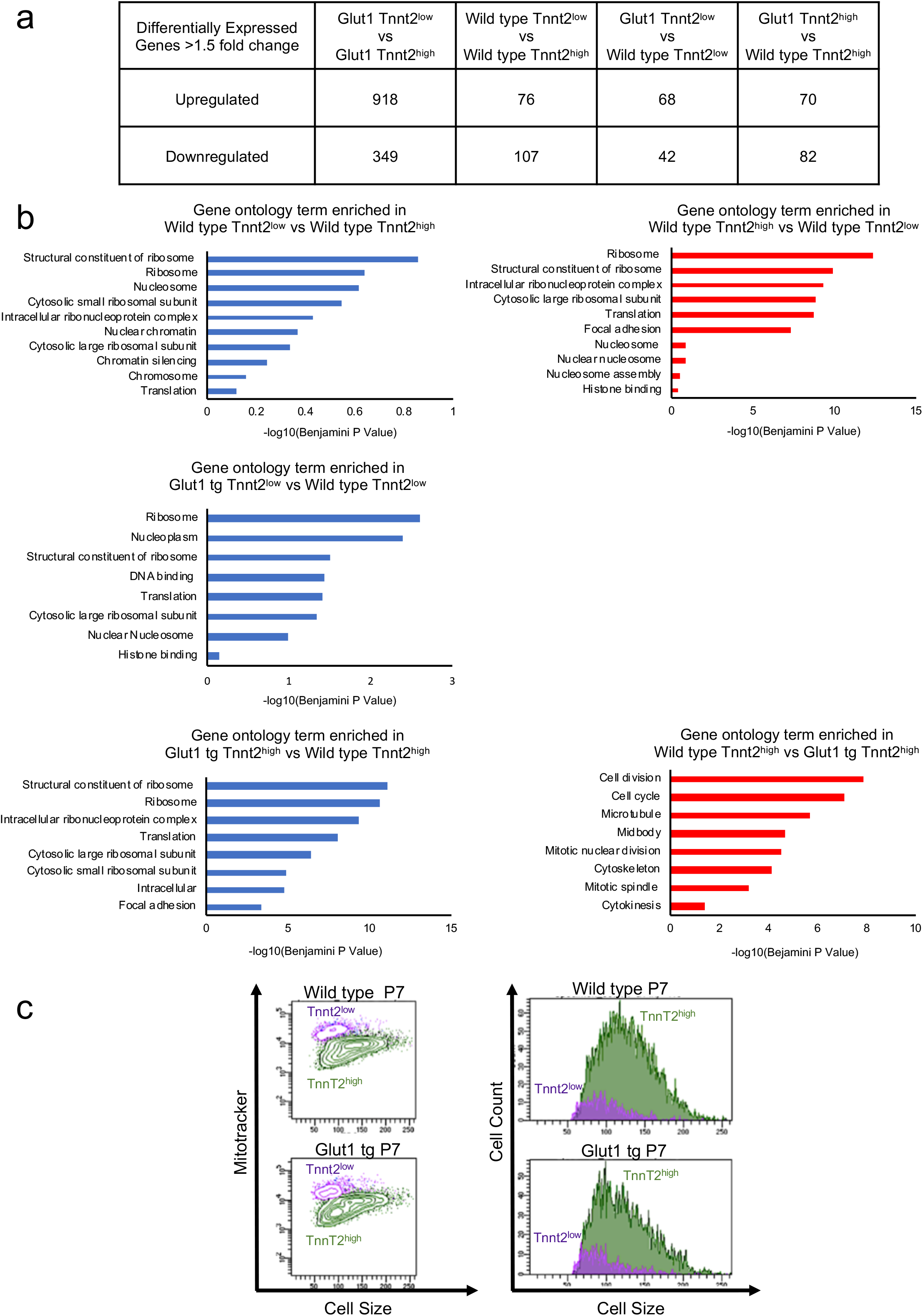
RNA-seq analysis of Tnnt2^high^ and Tnnt2^low^ cardiomyocytes from Wild type and Glut1 transgenic hearts at P1. a. 2-way comparison of Tnnt2^high^ and Tnnt2^low^ cardiomyocytes from Wild type and Glut1 transgenic hearts. Note the significant increase in the number of differentially expressed genes (DEGs) between Glut1 tg Tnnt2^high^ and Glut1 tg Tnnt2^low^ cardiomyocytes. b. Gene Ontology of DEGs among 4 populations: ^1^Glut1 tg Tnnt2^low^ vs Glut1 tg Tnnt2^high^; ^2^Wild type Tnnt2^low^ vs Wild type Tnnt2^high^; ^3^Glut1 tg Tnnt2^low^ vs Wild type Tnnt2^low^; ^4^Glut1 tg Tnnt2^high^ vs Wild type Tnnt2^high^. c. Histogram of the forward scatter of the 4 populations. Note that Tnnt2^low^ cardiomyocytes are smaller in size in both Wild type and Glut1 transgenic hearts.

**Supplemental Figure 5.**
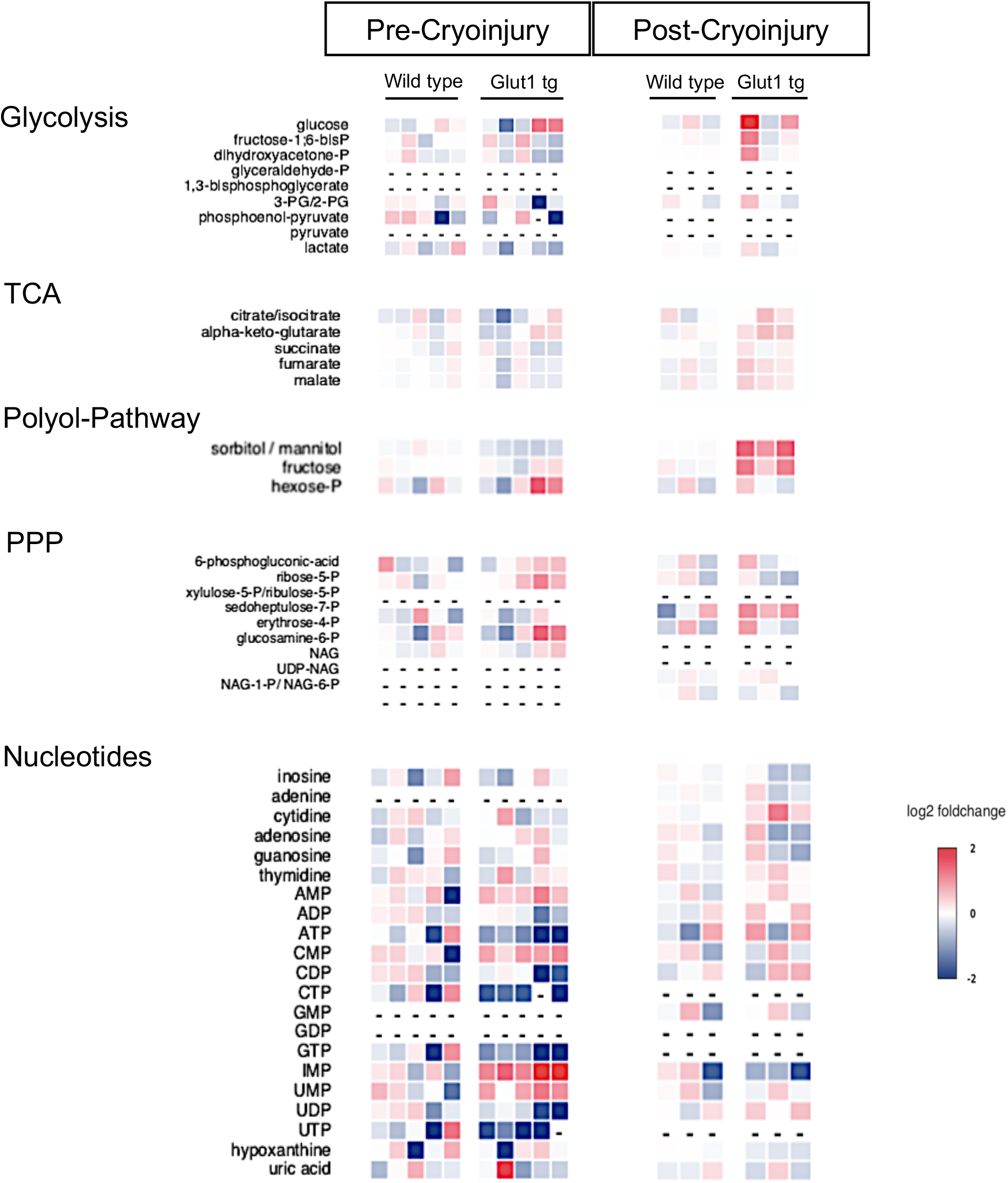
Neonatal cardiac metabolite differences prior and post-surgery. Heatmap of metabolites before and after surgery at P2 stage comparing wild type control hearts vs Glut1 transgenic hearts. Both groups were compared pre-injury and post-injury separately.

